# Partitioning covariance into causal and spurious components

**DOI:** 10.64898/2026.09.29.755348

**Authors:** Jonathan Henshaw

## Abstract

Statistical associations between two variables can arise via various causal mechanisms. For example, one variable may influence the other, or they may share common causes. Surprisingly little attention has been devoted to quantifying how such causal pathways contribute to the overall association between variables. Here, I present two methods for partitioning a covariance between variables *X* and *Z* into a causal component arising from the effect of *X* and *Z* and a spurious component arising via other pathways. Both methods rely on averaging causal decompositions over the distributions of the variables, but the decomposition and averaging are performed in different ways. The ‘gradient’ definition decomposes causal effects in the neighbourhood of naturally occurring values, making it relatively straightforward to estimate. It assumes that the cause variable *X* is normally distributed. In contrast, the ‘two-step’ definition decomposes large (‘step-wise’) changes in values and makes no distributional assumptions. I describe applications of these methods to analysing natural selection and illustrate them using the covariance between body size and the number of eggs carried by pregnant male dusky pipefish, *Syngnathus floridae*. I partition this total covariance into a causal component due to the effect of body size on egg number and a spurious component arising from variation in body size and egg number among collection sites.

---

Covariances quantify the extent of association between two variables. A robust non-zero covariance between variables *X* and *Z* – i.e., one that persists even in sufficiently large random samples – is evidence of a causal link between these variables. We can broadly classify such causal links into three non-mutually-exclusive types^1^:

1. *X* influences *Z* (*X* → *Z*)
2. *Z* influences *X* (*Z* → *X*),
3. both *X* and *Z* are influenced by some underlying third variable(s) (*X* ← *C* → *Z*).

The arrows above should be understood to allow for both direct and indirect causal effects (e.g., *X* → *Z* also includes causal chains of the form *X* → *⋯* → *Z*). Importantly, the mere presence of covariance is insufficient to determine the nature of the underlying causal links: this is the insight behind the saying ‘correlation does not imply causation’. However, even if the causal relationships between *X* and *Z* are completely understood, we may still ask: *How much of the covariance between X and Z arises from causal effects of X on Z, and how much via other causal pathways?* In the context of linear and additive effects, this question was fully resolved in the 1930s by Sewall Wright’s method of path coefficients (often known as ‘path analysis’) (Wright 1934; Loehlin 2004). In a fully general setting, allowing for arbitrary relationships among variables, this question has received surprisingly little attention: I am aware of only the pioneering study by Zhang and Bareinboim (2018) (see also Plečko and Bareinboim 2023). Here, I outline two potential solutions.

My own motivation for this problem is rooted in evolutionary biology. I consequently begin by briefly outlining some of the many applications of covariances in evolutionary theory, as well as some existing approaches for analysing their causal determinants. Readers from other fields who do not need this evolutionary context may wish to skip to the following section (‘A toy scenario’).

Unsurprisingly for such a fundamental statistical construct, covariances show up everywhere in the evolutionary literature. As in other fields, they serve as straightforward and familiar summary statistics for the linear association between two variables. Covariances also arise in a particularly natural way in evolutionary theory, for example in the context of predicting evolutionary change or measuring genetical associations between traits (e.g., Wright 1934; Price 1970, 1972; Lande and Arnold 1983; Moore et al. 1997; Arnold et al. 2008; Walsh and Lynch 2018; Svensson et al. 2021). A particularly central example is the *selection differential*, which quantifies the difference in mean trait values between the population as a whole and the parents of offspring, where the latter are weighted by their individual fitness (Walsh and Lynch 2018). (For the purposes of our current discussion, we can think of ‘fitness’ as the number of offspring an individual produces over their lifetime, although more complex formulations are sometimes necessary.) In the hypothetical scenario where parents pass on their trait values to offspring with perfect fidelity, the selection differential represents the difference in trait values between parents and offspring, assuming a constant environment. Importantly, the selection differential can be written as a covariance *S* = cov(*X, w*), where *X* represents individual trait values and *w* stands for individual relative fitness (‘relative’ means that fitness has been normalized by dividing each value by the population mean fitness) (Lande and Arnold 1983).

Wright (1920, 1921) recognized early on that selection can arise via multiple causal pathways and developed the method of path analysis to quantify the relative importance of these pathways. When effects are linear and additive, path analysis allows the selection differential to be expressed as the sum of effects via each contributing causal pathway (Wright 1934; Loehlin 2004). Wright’s framework was later extended to study non-linear selection by including quadratic and interaction terms (Scheiner et al. 2000) and subsequently generalised to allow for arbitrary functional relationships between effects and their causes (Morrissey 2014, 2015; Henshaw et al. 2020). This work culminates in an explicit expression 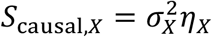 for the component of the selection differential that is due to direct or indirect causal effects of the focal trait on fitness. Here, 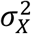 is the variance in trait values and *η*_*X*_ is the *extended selection gradient*, defined as the mean expected rate of change in individual fitness predicted if an experimenter intervened to slightly increase a random individual’s trait value *X* (Morrissey 2014, 2015; Henshaw et al. 2020). This construction is closely related to the ‘gradient’ definition of causal covariance presented below (see also Lehtonen and Otsuka 2023; Frank 2026). It represents one strand of a small but growing literature that applies modern causal inference theory (Pearl 2009; Hernán and Robins 2020) to questions in evolutionary biology (e.g., von Hardenberg and Gonzalez-Voyer 2013; Okasha and Otsuka 2020; Queller 2020; von Hardenberg and Gonzalez-Voyer 2025; Ringen et al. 2026).

## A toy scenario

Although my motivation is evolutionary, I will first present the definitions in a general setting. To keep things conceptually and notationally manageable, I will focus on the simplest possible causal structure with which we can approach the question. Most of the ideas developed in this setting transfer readily to more complex causal structures (e.g., to those that can be represented by directed acyclic graphs: Pearl 2009). I will also stick to the standard linear notion of covariance, although various non-linear covariance concepts could be approached analogously.

Our set-up is encoded by the path diagram in Figure 1a. A random variable *Z* is fully determined by random variables *X* and *Y*. We allow that *X* and *Y* may covary due to unspecified common causes. Note, in particular, that causal influences only go in one direction in this scenario: I am excluding the possibility that *Z* influences *X* or *Y*. For simplicity, suppose that we have full knowledge of:

**Figure 1:**
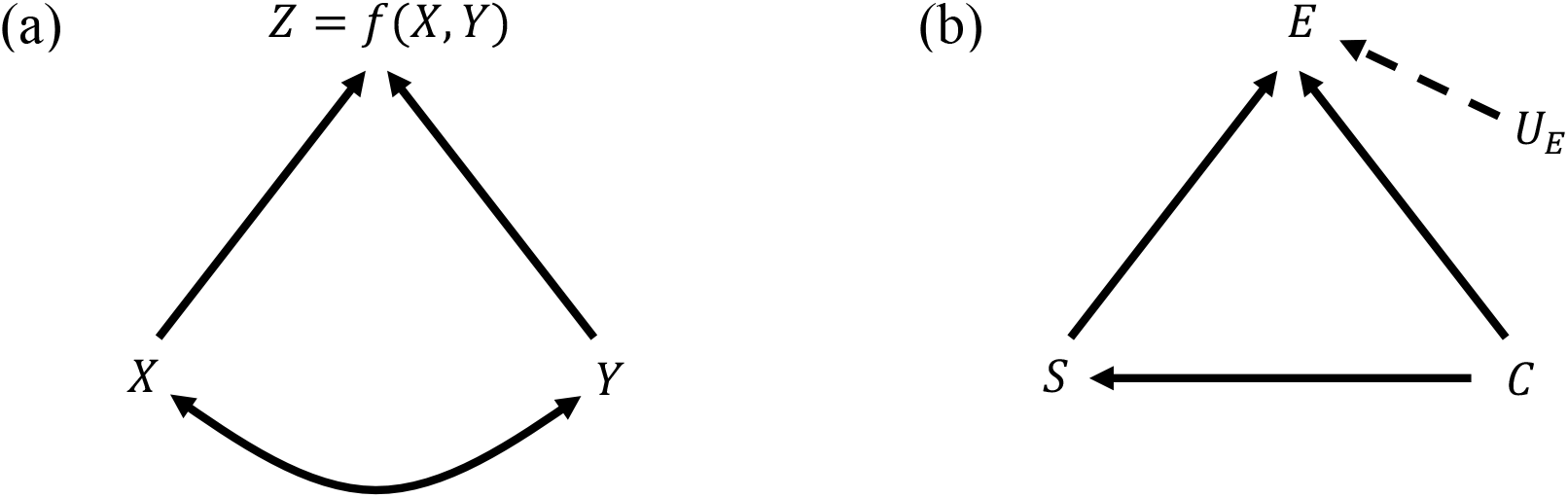
**(a)** A simple causal scenario. The random variable *Z* is fully determined by *X* and *Y* (single-headed arrows). The variables *X* and *Y* may covary due to unspecified shared causes (double-headed arrow). **(b)** The assumed causal relationships among body size *S*, the number of eggs carried *E*, and the collection event *C*, in the worked example on pregnant male dusky pipefish. The background variable *U*_*E*_ represents unmeasured sources of variation in egg number and is assumed to be independent of (*S, C*).

1. the joint probability distribution of *X* and *Y*, and
2. the functional relationship *Z* = *f*(*X, Y*) between *Z* and its causal determinants.

In practice, this information could be estimated using the empirical distribution of *X* and *Y* and an appropriate regression of *Z* on these variables. We then ask: How much of cov(*X, Z*) arises via the *causal* pathway *X* → *Z*, and how much via the *spurious* pathway *X* ↔ *Y* → *Z*? Can we partition the covariance cov(*X, Z*) into a causal component cov_casual_(*X, Z*) and a spurious component cov_spurious_(*X, Z*) such that

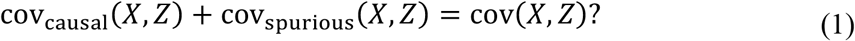

## Minimal requirements

Before we begin, it is helpful to outline some minimal requirements that any candidate solution should satisfy. First, suppose that *X* is independent of *Y*. In this case, any covariance between *X* and *Z* cannot arise spuriously via covariation between *X* and *Y*. Hence, our first requirement is:

(R1) *Independence:* If *X ⊥ Y* then cov_casual_ (*X, Z*) = cov(*X, Z*) and cov_spurious_(*X, Z*) = 0.

Second, suppose the effects of *X* and *Y* on *Z* are *additive*, meaning that there exist functions *f*_*X*_ and *f*_*Y*_ such that

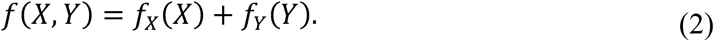

In this case, we have:

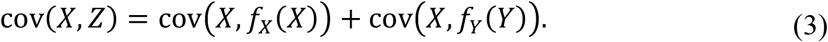

The first term here captures covariance arising via the effect *X* → *Z*, while the second term captures covariance arising via the effect *Y* → *Z* and the covariance between *X* and *Y*. The causal partitioning of cov(*X, Z*) is thus straightforward in this additive case, which leads to our second requirement:

(R2) *Additivity:* If *Z* = *f*_*X*_(*X*) + *f*_*Y*_(*Y*) then cov_casual_ (*X, Z*) = cov=*X, f*_*X*_(*X*)) and cov_spurious_ (*X, Z*) = cov=(*X, f*_*Y*_(*Y*)).

Lastly, suppose that the function *f*(*⋅,⋅*) is symmetrical in its arguments and that *X* ≡ *Y*. Then the causal effect of *X* on *Z* is identical in form to the association of *X* with *Z* via *Y*. The causal and spurious pathways linking *X* to *Z* should consequently be equal in magnitude:

(R3) *Symmetry:* If *X* ≡ *Y* and *f*(*x, y*) = *f*(*y, x*) for all (*x, y*), then 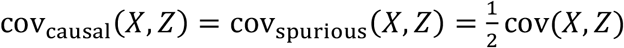.

Importantly, these three requirements do not uniquely determine a solution to the problem in equation (1). I will consider two distinct constructions of the causal and spurious covariances that do fulfill these three requirements.

## Two-step definition

For the first construction, it is helpful to start with a simpler question. Imagine that, via some external intervention, we change the value of the ordered pair (*X, Y*) from (*x*_1_, *y*_1_) to (*x*_2_, *y*_2_). As a result of this intervention, the value of *Z* changes from *z*_1_ = *f*(*x*_1_, *y*_1_) to *z*_2_ = *f*(*x*_2_, *y*_2_). How much of this change in *Z* is due to us intervening on *X* and how much is due to us intervening on *Y*? To approach this question, we could consider changing the values of *X* and *Y* sequentially, rather than simultaneously. The change in *X* while holding *Y* constant would provide an estimate of the causal effect of *X* alone. Here we already encounter a problem, however: it can make a difference in which order we alter these variables (Figure 2; cf. Pearl 2009; Singal et al. 2021; Singal and Michailidis 2024).

**Figure 2:**
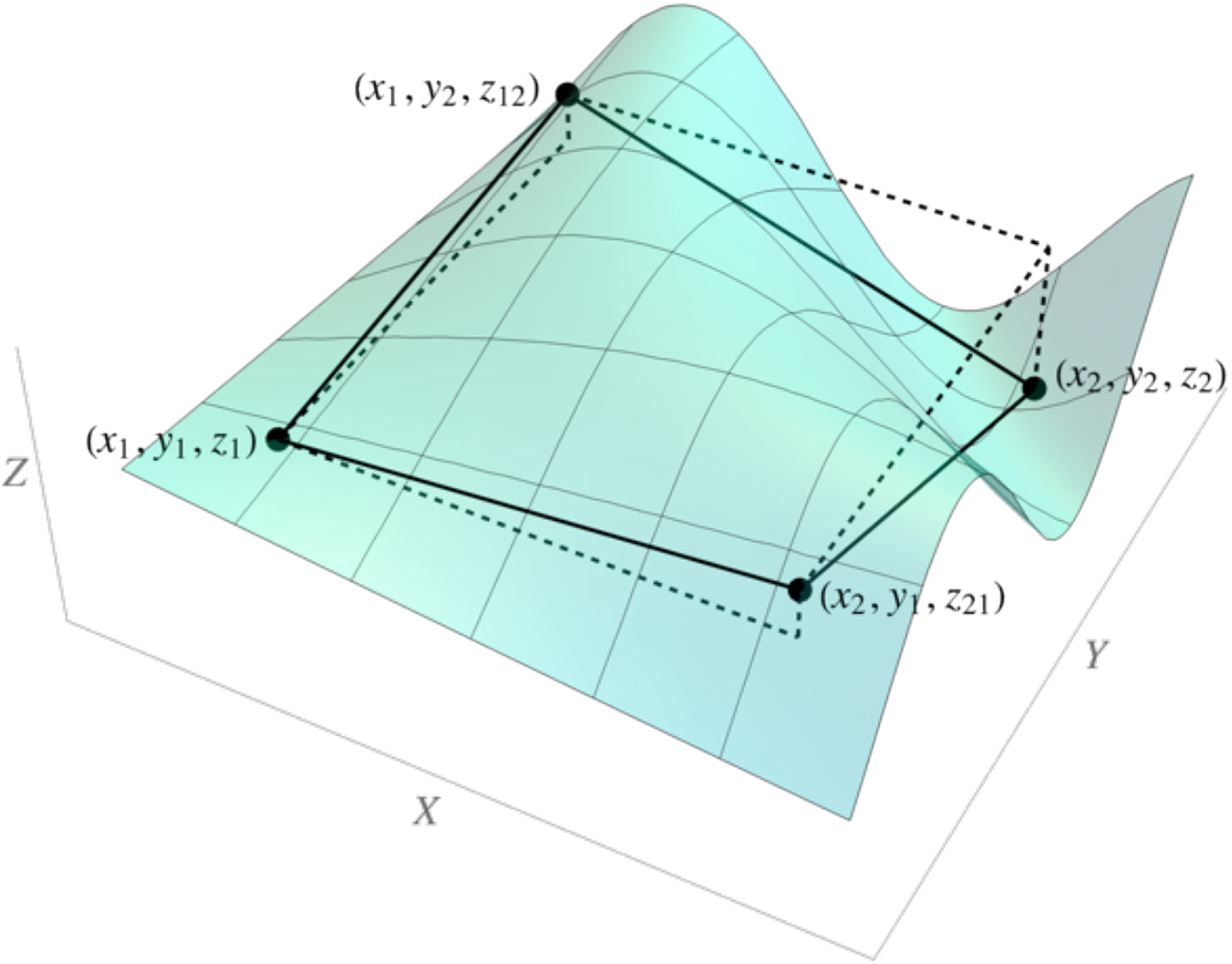
A construction involved in the two-step definition of causal and spurious covariances. If we change (*X, Y*) from (*x*_1_, *y*_1_) to (*x*_2_, *y*_2_), the value of *Z* changes by *z*_2_− *z*_1_. We can attribute this difference to the effects of *X* and *Y* by considering changes in each variable sequentially. However, it makes a difference in which order we change *X* and *Y*. In the example depicted, even the sign of the imputed effects differs: If we change *X* first, then the effect *z*_12_ – *z*_1_ of *X* is positive and the effect *z*_2_ – *z*_12_ of *Y* is negative. If we change *Y* first, then the effect *z*_2_− *z*_21_ of *X* is negative and the effect *z*_21_ – *z*_1_ of *Y* is positive. The two-step definition effectively takes the average of these two imputed effects over pairs (*x*_1_, *y*_1_) and (*x*_2_, *y*_2_) drawn independently from the joint distribution of (*X, Y*). Note that if the effects of *X* and *Y* on *Z* are additive, then the four labelled points are coplanar and the imputed effects of *X* and *Y* are the same no matter which order we change the variables.

Suppose we first change *X* from *x*_1_ to *x*_2_, measure the change in *Z*, and then change *Y* from *y*_1_ to *y*_2_. We then obtain the following causal breakdown of the difference *z*_2_ – *z*_1_:

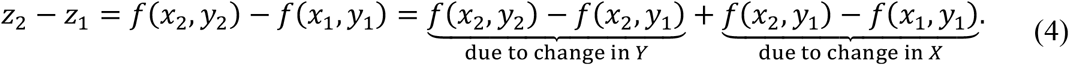

I refer to this as the *X*-*first* or *causal-first* breakdown, where the latter term assumes that we are interested in the causal effect of *X* on *Z*. On the other hand, if we first change *Y* and then *X*, we obtain:

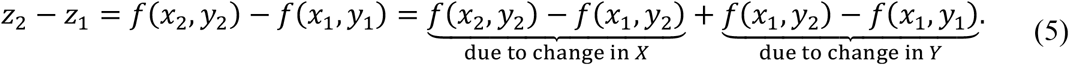

This is the *Y*-*first* or *spurious-first* breakdown. If the effects of *X* and *Y* on *Z* are non-additive, then these two breakdowns may result in different estimates of how changes in *X* and *Y* contribute to change in *Z* (Figure 2). As we will see, however, there is a natural causal partitioning of the covariance that does not depend on whether we rely on the causal-first or spurious-first breakdowns.

Let’s now take a short excursion. Instead of focussing on fixed values of *X* and *Y*, consider independently and identically distributed ordered pairs (*X, Y*) and (*X*\*, *Y*\*). In other words, we create an independent copy of the joint distribution of *X* and *Y*. Corresponding to these are independent variables *Z* = *f*(*X, Y*) and *Z*\* = *f*(*X*\*, *Y*\*). The covariance between *X* and *Z* can then be written as:

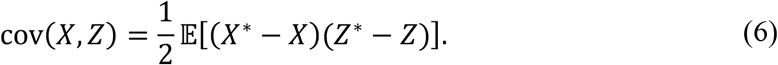

This ‘independent copy’ formula can be interpreted as follows: We repeatedly draw independent pairs of values from the joint distribution of (*X, Y*) (i.e., pairs of the form =((*x*_1_, *y*_1_), (*x*_2_, *y*_2_))). For each such pair, we calculate *z*_/_ = *f*(*x*_1_, *y*_1_) and *z*_2_ = *f*(*x*_2_, *y*_2_) and multiply the difference in *X*-values by the corresponding difference in *Z*-values. The covariance between *X* and *Z* is equal to *one half* of the mean of such products of differences. By substituting in either the causal-first or the spurious-first breakdowns from equations (4) or (5) – it makes no difference which one we choose – we can express the factor *Z*\* − *Z* in terms of changes due to differences in *X* and changes due to differences in *Y*. We thereby obtain the *two-step definition* of causal and spurious covariances:

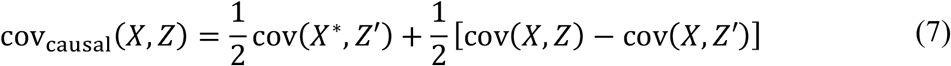

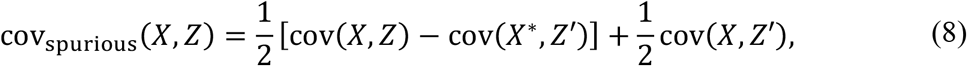

where *Z*′ = *f*(*X*\*, *Y*). These equations may appear cumbersome initially, but they have straightforward interpretations. Under natural conditions (i.e. in the absence of any external intervention), *X* may covary with *Z* due to both the causal effect *X* → *Z* and the spurious relationship *X* ↔ *Y* → *Z*. The creation of a new variable *X*\* that is independent of (*X, Y*) allows these two effects to be separated (Figure 3). Since *X*\* and *Y* are independent, the covariance between *X*\* and *Z*′ = *f*(*X*\*, *Y*) is determined entirely by the causal effect *X*\* → *Z*′. On the other hand, the covariance between *X* and *Z*′ is determined entirely by the spurious relationship *X* ↔ *Y* → *Z*′. We thus obtain two distinct ways of conceptualizing the causal and spurious components of the covariance between *X* and *Z*:

**Figure 3:**
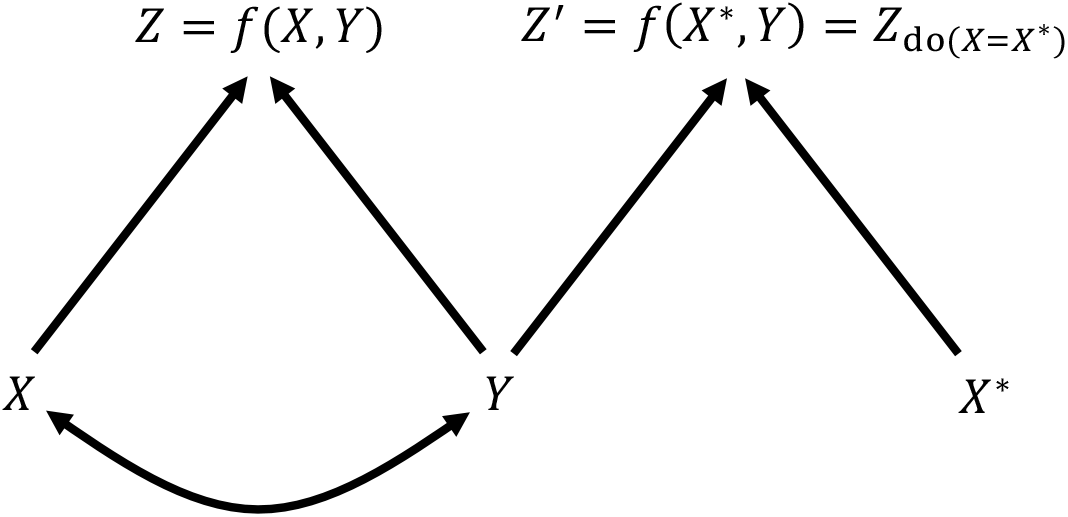
The causal structure underlying the two-step definition of causal and spurious covariances. A new variable *X*\* is introduced that is identically distributed to *X* but independent of (*X, Y*). The variable *Z*^′^ = *f*(*X*\*, *Y*) is determined by the new *X*\* and the original *Y* according to the same functional relationship as the original *Z*. Since *X*\* is independent of *Y*, we might take cov(*X*\*, *Z*^′^) to represent the causal covariance between *X* and *Z*, with the remaining covariance being spurious (D1). Alternatively, since *X* has no causal effect on *Z*′, but only a spurious relationship via *Y*, we might take cov(*X, Z*^′^) to represent the spurious covariance between *X* and *Z*, with any remaining covariance being causal (D2). The two-step definition simply averages the definitions (D1) and (D2).

(D1) Define the causal covariance as any covariance remaining after ‘breaking’ the spurious relationship between *X* and *Y*. The causal covariance is then cov(*X*\*, *Z*^′^) and the spurious covariance is the remainder cov(*X, Z*) − cov(*X*\*, *Z*^′^).

(D2) Define the spurious covariance as any covariance remaining after ‘breaking’ the causal effect of *X* on *Z*. The spurious covariance is then cov(*X, Z*^′^) and the causal covariance is the remainder cov(*X, Z*) − cov(*X, Z*^′^).^2^

There is no apparent reason to favour one viewpoint over the other. It is consequently satisfying that the two-step definition simply averages the quantities arising from definitions (D1) and (D2). Note, moreover, that the two-step definition is identical in form regardless of whether we substitute the causal-first or the spurious-first breakdowns (equations (4) or (5) respectively) into equation (6). Intuitively, this is because each pair of values (*x*_1_, *z*_1_) and (*x*_2_, *z*_2_) is accounted for twice under the expectation in equation (6), but in different roles: in one instance, we have (*X, Z*) = (*x*_1_, *z*_1_) and (*X*\*, *Z*\*) = (*x*_2_, *z*_2_), and in the other, (*X, Z*) = (*x*_2_, *z*_2_) and (*X*\*, *Z*\*) = (*x*_1_, *z*_1_). Consequently, regardless of which breakdown we work with, the effect on *Z* of the change from *x*_1_ to *x*_2_ is quantified once against the background of *y*_1_ and once against the background of *y*_2_.

The two-step definition can be extended readily to more general settings. Following Pearl (2009), we write do(X = *x*) for an external intervention that sets the value of *X* to *x* without regard for how *X* would normally be determined. The random variable *Z*_do(*x=x)*_ is obtained by intervening to set *X* equal to *x* and then observing *Z*. Using this notation, we can write 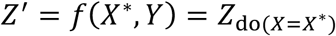. Consequently, if *X*\* is independent of (*X, Y*) but follows the same distribution as *X*, then we can rewrite the two-step definition as:

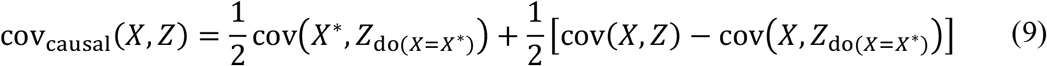

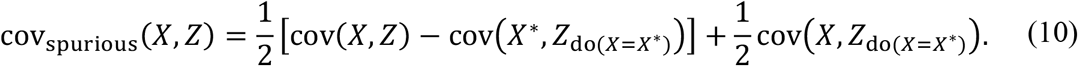

This definition extends to any causal structure that can be represented by a directed acyclic graph. A range of techniques exists for estimating the distribution of 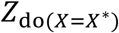, including the useful *backdoor criterion*, which formalises the art of ‘controlling for’ confounders when estimating the causal effect of a predictor (for details see Pearl 2009).

## Gradient definition

The two-step definition analyses differences in *Z* corresponding to pairs of values drawn from the joint distribution of (*X, Y*) (i.e. pairs of the form =((*x*_1_, *y*_1_), (*x*_2_, *y*_2_))). Since each member of a pair is drawn independently, the differences in the analysed *X*- and *Y*-values (i.e., the differences *x*_2_ – *x*_1_ and *y*_2_ – *y*_1_) may be large. In contrast, I next present a construction based on infinitesimal changes in the focal variable *X*. Importantly, this analysis assumes that *X* is normally distributed.

Our starting point is Stein’s lemma, which provides a link between the covariance cov(*X, Z*) and the mean derivative of the conditional expectation E (*Z*|*X*) with respect to *X* (Stein 1981; Lande and Arnold 1983; Liu 1994):

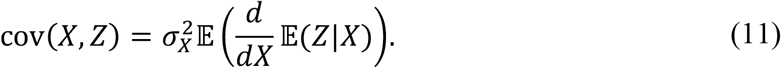

Here 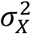 is the variance of *X*. The expression 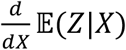 quantifies how small changes in *X* are associated with changes in *Z*, whether due to causal effects of *X* and *Z* or spurious associations via confounding variables. Note that Stein’s lemma assumes the normality of *X*. Now, by the law of total expectation, we have:

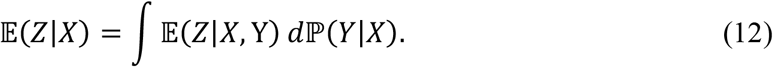

Substituting this expression into equation (11), we can differentiate under the integral sign and apply the product rule of calculus to obtain (Henshaw et al. 2020):

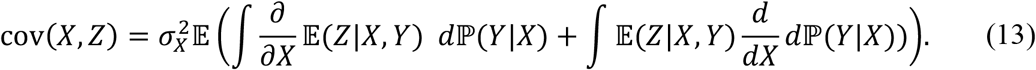

We can further simplify the first term inside the brackets by integrating under the derivative and applying the law of total expectation:

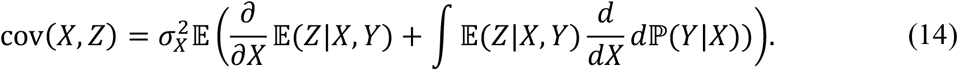

The first term describes the average effect on *Z* of infinitessimal changes in *X*, while holding *Y* constant. It thus quantifies the average causal effect of *X* on *Z*. This leads to the *gradient definition* of the causal covariance:

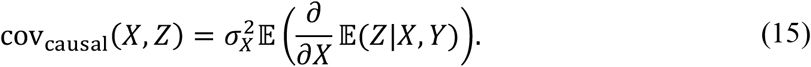

Note that in our toy scenario, in which *Z* is fully determined by *X* and *Y*, we have E (*Z*|*X, Y*) = *f*(*X, Y*) = *Z*. Accounting for additional sources of variation in *Z* is straightforward, however (see below). In the special case where *X* is a trait value and *Z* = *w* is relative fitness, cov_casual_(*X, Z*) equals the extended selection gradient on *X* (Morrissey 2014, 2015; Henshaw et al. 2020).

The second term in equation (14) describes the average effect on *Z* of changes in the distribution of *Y* that are associated with (but not caused by) infinitesimal changes in *X*. The gradient definition of the spurious covariance is then:

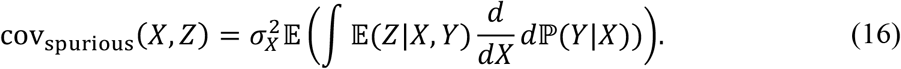

In the special case where *X* is a trait value and *Z* = *w* is relative fitness, cov_spurious_(*X, Z*) quantifies selection on the trait *X* that does not arise from the causal effect of *X* on fitness, but rather due to its covariation with *Y* and the effect of *Y* on fitness.

Like the two-step definition above, the gradient definition can be formulated in a more general way that accommodates more complex causal structures. We define the *causal derivative* of *Z* with respect to *X* as the rate at which *Z* changes under an infinitesimal intervention on *X* (Henshaw et al. 2020):

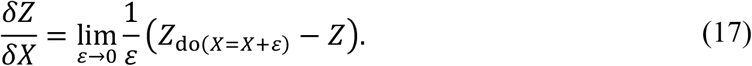

Here, the intervention increases the natural value of *X* by a small amount *ε*. Under the causal assumptions encoded in Figure 1a, we have 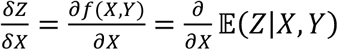. The gradient definition can then be rewritten as:

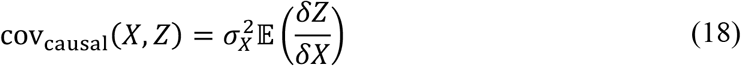

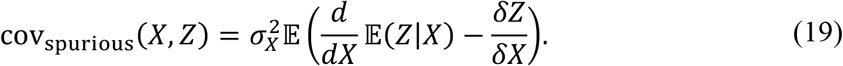

This definition can be applied to any causal structure that can be represented by a directed acyclic graph, provided that the random variable *X* is normally distributed and that the causal derivative exists. The latter assumption will not always hold; in particular, it excludes cases where *Z* is a discrete random variable. In such cases, we can often get around the issue by considering the conditional expectation of *Z* averaged over some unmeasured background noise. More concretely, suppose first that ***B*** is a *backdoor set* for the effect of *X* on *Z*, meaning that (i) ***B*** contains no descendants of *X*, and (ii) ***B*** blocks all backdoor (i.e., non-causal) paths from *X* to *Z* (Pearl 2009; Henshaw et al. 2020). Then we can replace 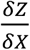 with 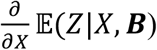 in equations (18) and (19), thereby averaging over noise in *Z* that is not relevant to calculating the causal effect of *X* on *Z*. In many cases, the expectation E (*Z*|*X*, ***B***) will be a continuous random variable even when *Z* itself is not.

### Worked example: Effect of body size on reproductive success in male pipefish

To illustrate these two causal breakdowns of covariance, I analysed the effect of body size on reproductive success in pregnant male dusky pipefish, *Syngnathus floridae*, using data collected by Kenyon Mobley and colleagues on the Eastern coast of North America (Mobley and Jones 2007, 2009, 2013; Mobley et al. 2014). The authors collected pregnant males from five sites, one of which was sampled in two separate years, resulting in six separate collection events *C*. They measured each male’s body size *S* and counted the number of eggs he was carrying *E*. The data set and R code for the worked example are provided as Supplementary Materials.

I assume that the variables can be represented by the causal diagram *C* → *S* → *E* with *C* → *E* (Fig. 1b). In other words, egg number depends on size and both size and egg number can depend on collection event. Remaining variation in egg number is assumed to result from an unmeasured background variable *U*_*E*_ that is independent of (*S, C*). Note that the following analyses would remain unchanged if we replaced *C* → *S* with *C* ↔ *S* in the causal diagram or allowed for both effects simultaneously. Under these assumptions, *C* forms a backdoor set for the effect *S* → *E*, and so controlling for *C* allows us to isolate the effect of size on egg number. I modelled this relationship using multiple linear regression, with size *S*, collection event *C*, and their interaction as predictors and egg number *E* as the response variable (Fig. 4). I then estimated the causal and spurious covariances of size on egg number using both the two-step and gradient definitions. Since a male’s fitness depends not only on the number of eggs he carries in a given season, but also on several other variables (such as the male’s reproductive lifespan, the quality of the eggs, etc.), the covariance between body size and egg number is not strictly speaking a selection differential. Nonetheless, egg number is an important component of fitness and so analysing its association with body size sheds light on one portion of the causal network that determines fitness.

**Figure 4:**
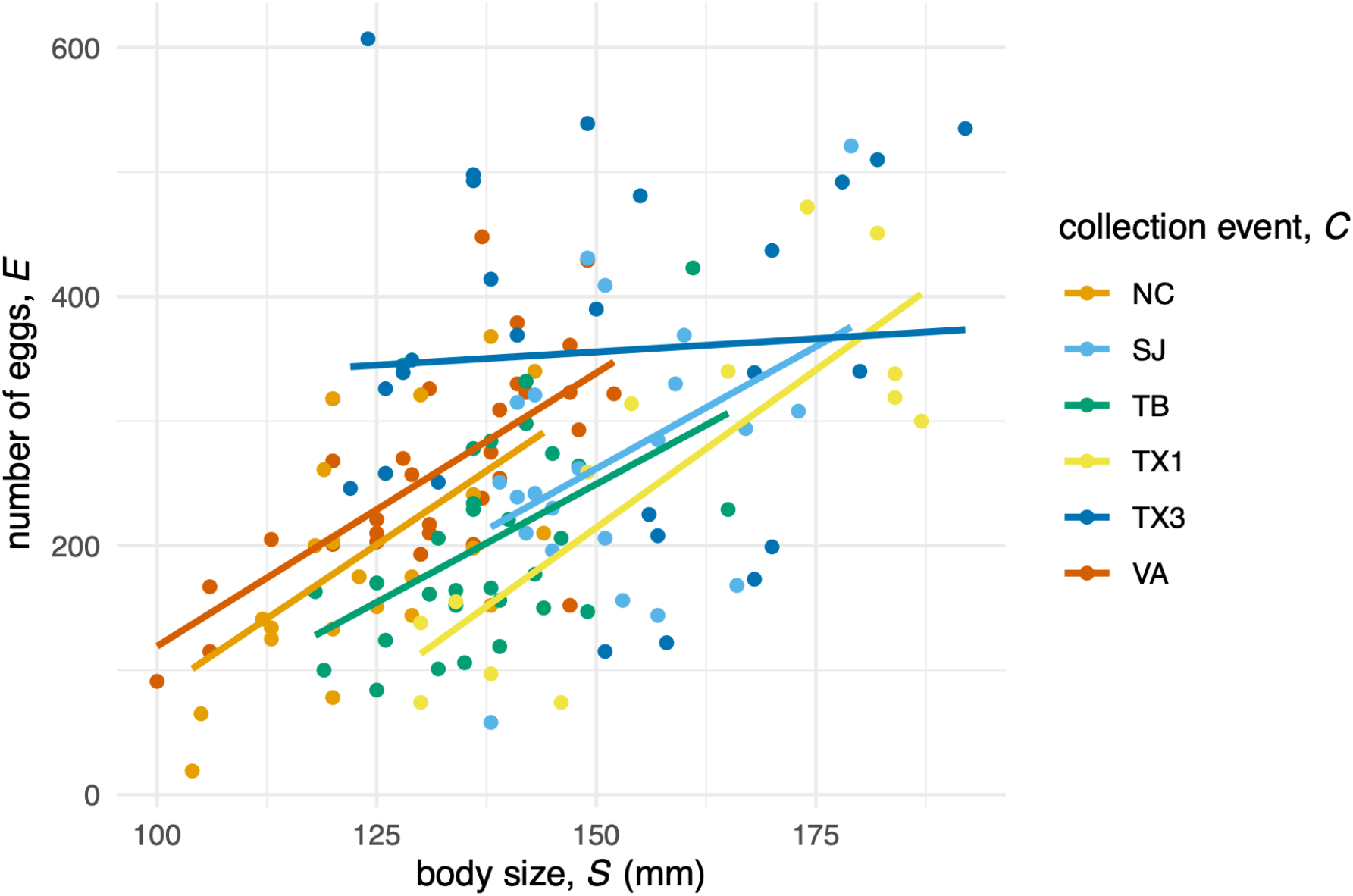
The predicted relationship between effects of body size *S* and collection event *C* on the number of eggs carried *E* in the worked example on pregnant male dusky pipefish (data from Mobley and Jones 2007, 2009, 2013; Mobley et al. 2014). Regression lines are based on a linear regression model including size, collection event and their interaction as predictors.

For the two-step definition, I drew *N* = 10^6^ independent pairs of samples (*S, C*) and (*S*\*, *C*\*) from the empirical joint distribution of the predictors. I then used the regression model to simulate *E*|*S, C* and *E*|*S*\*,*C* for each sampled pair. Lastly, I substituted these values into equations (7) and (8) to obtain the causal and spurious covariances. For comparison, I also estimated the total covariance cov(*S, E*) based on the re-sampled (*S, C*) values and the model-simulated values of *E*|*S, C*.

For the gradient definition, I used the regression model to estimate E (*E*|*S, C*) and E (*E*|*S* = *S* + *ε, C*), with *ε* = 0.1, for each observed set of predictors (*S, C*). Following equation (17), I then estimated the causal gradient for each set of predictors numerically as:

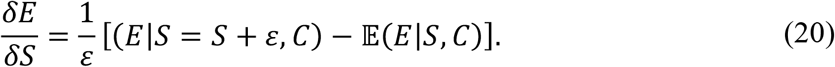

Finally, I averaged 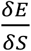 across all observations and multiplied the result by the variance 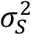 to obtain the causal covariance, following equation (18). The spurious covariance was calculated by subtracting this result from the total covariance cov(*S, E*). In this simple model, the causal gradient 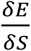 quantifies the effect of size on egg number while holding collection event constant. It consequently equals the collection-event-specific slopes in the regression model. The mean causal gradient 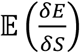 simply averages these regression slopes across all collection events, weighted by the number of individuals sampled in each collection event.

The two causal breakdowns are quantitatively similar (Table 1). Both the total covariance cov(*S, E*) and the causal covariance cov_casual_(*S, E*) are positive, but the causal covariance is larger, suggesting that the causal effect of body size on egg number is stronger than the statistical association between these variables might suggest. In line with this, the spurious covariance cov_spurious_(S, *E*) is negative, suggesting that the non-causal pathway *S* ← *C* → *E* reduces the statistical association between *S* and *E*.

**Table 1.** Comparison of causal breakdowns of the covariance of body size *S* with egg number *E* in dusky pipefish (data from Mobley and Jones 2007, 2009, 2013; Mobley et al. 2014). Correlation-like quantities *ρ* in brackets are calculated by dividing the covariances by *σ*_*S*_*σ*_*E*_. Note that the total covariance cov(*S, E*) presented for the gradient definition was calculated directly from the empirical distribution of (*S, E*), whereas the total covariance for the two-step definition was derived by re-sampling (*S, C*) and simulating *E*|*S, C* under the regression model.

|  | two-step definition | gradient definition |
| --- | --- | --- |
| $\text{cov}(S, E)$ | 1043.2 ($\rho = 0.48$) | 1049.9 ($\rho = 0.49$) |
| $\text{cov}_{\text{causal}}(S, E)$ | 1189.8 ($\rho = 0.55$) | 1229.3 ($\rho = 0.57$) |
| $\text{cov}_{\text{spurious}}(S, E)$ | -146.5 ($\rho = -0.07$) | -179.4 ($\rho = -0.08$) |

## Comparison of the two definitions

Here I discuss some similarities and differences between the two-step and gradient definitions of causal and spurious covariances. Broadly speaking, both definitions of causal covariance are based on an average causal effect, but the way in which the average is taken differs between the definitions. For the two-step definition, the average is taken over randomly chosen large differences in both variables – the focal cause and effect – while respecting the structural relationships between the variables. In contrast, the gradient definition deals with infinitesimal changes in the focal cause. Neither approach is fundamentally more valid than the other, as they answer different questions.

It is straightforward to verify that both definitions satisfy the proposed minimal requirements relating to independence, additivity and symmetry ((R1) through (R3) above). The symmetry property, in particular, sets these definitions apart from the previous definition of Zhang and Bareinboim (2018). Their definition is equivalent to (D2) if one averages all estimates over the empirical distribution of the variables (as suggested in their footnote 8). Neither (D1) nor (D2) fulfils the symmetry requirement on its own, although both definitions satisfy the independence and additivity requirements. In contrast, the two-step definition ensures symmetry by averaging the quantities defined by (D1) and (D2). Further, the definitions (D1) and (D2) are based on two distinct mechanisms for isolating causal effects (Figure 3): either eliminate all spurious relationships (D1) or disrupt the causal effect of the focal variable (D2). There is no obvious reason to favour one approach over the other. This might be seen as an additional argument in favour of the two-step definition, which naturally accounts for both mechanisms, over definitions that do not.

Importantly, unlike the overall covariance, neither definition of causal covariance is symmetric in its arguments (i.e., cov_casual_(*X, Z*) ≠ cov_casual_(Z, X)). This is to be expected, as associations are symmetric, whereas causal effects are not. Indeed, if the causal relationships between *X* and *Z* can be described using a directed acyclic graph (i.e., without any feedback loops), then at least one of these causal covariances must equal zero.

The two definitions also differ in their distributional assumptions. The gradient definition applies only when the focal variable *X* is normal in distribution (the other variables may follow any probability distribution). Extensions of Stein’s lemma to non-normal variables do exist (Goldstein and Reinert 1997, 2005), but apparently do not allow for clean causal interpretations (see Appendix). Even when the focal variable *X* is not normal, the mean causal derivative 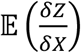 has a clear causal interpretation as the average rate of change in *Z* under infinitesimal interventions on *X*. However, the straightforward link between 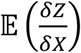 and the covariance cov(*X, Z*) is lost for non-normal *X*.

In contrast, the two-step definition can be applied to focal variables following any probability distribution. However, an unintuitive property of this definition is that it considers causal effects under conditions where the natural associations between *X* and its covariates have been disrupted (Figure 3). For example, if *X* and *Y* covary strongly, then there will be combinations of *X*- and *Y*-values that rarely or never occur under natural conditions. In contrast, *X*\* is independent of *Y* by construction and so the joint distribution of (*X*\*, *Y*) is simply the product of the marginal distributions. Estimating the effects of such unusual combinations on *Z* may be difficult or impossible in practice due to a paucity of data. In contrast, the gradient definition requires only that we can estimate causal effects in the neighbourhood of naturally occurring values in the joint distribution of (*X, Y*).

## Discussion

The ideas in this work are not fundamentally new. The causal gradient is closely related to the concept of ‘controlling for’ in statistics, which is made precise in the causal inference literature through the backdoor criterion (Pearl 2009). The average value of this gradient is often referred to as the *average partial effect* or the *average marginal effect*, especially in economics (Woolridge 2002). The relationship between covariances and average partial effects has been exploited in evolutionary theory for decades (Lande and Arnold 1983).

In the two-step definition, the counterfactual (D1) is closely related to the *average treatment effect* in the causal inference literature (Hernán and Robins 2020). Averaging over the alternative counterfactuals (D1) and (D2) is reminiscent of the Shapley value in game theory, which assigns a fair payout to each player in a cooperative group by averaging payouts over all orders in which the players could have joined the group (Shapley 1953; Osborne and Rubinstein 1994). Indeed, approaches based on the Shapley value have already been applied to decomposing causal effects (Singal et al. 2021; Jung et al. 2022; Singal and Michailidis 2024).

Averaging over alternative counterfactuals is a useful device when there are non-additive interactions between causal pathways (Singal and Michailidis 2024). An alternative approach is to explicitly separate additive effects from interactions (VanderWeele 2013; VanderWeele and Tchetgen Tchetgen 2014). For example, one could partition the total association between two variables into a causal component, a spurious component, and a component that arises from interactions between causal and spurious pathways. Interestingly, if we apply such a breakdown to the independent-copy formula in equation (6) (following VanderWeele and Tchetgen Tchetgen 2014), then the interaction component disappears when taking the expectation. This is because each set of values (*x*_1_, *x*_2_, *z*_1_, *z*_2_) appears equally often with (*X, Z*) = (*x*_1_, *z*_1_) and (*X*\*, *Z*\*) = (*x*_2_, *z*_2_) as vice versa, and so the interaction effect cancels out in the aggregate. The gradient formula sidesteps the issue of non-additivity by focussing on infinitesimal changes, where local linearity ensures that the contributions of different causal pathways are additive (contrary to the claim of Singal and Michailidis 2024).

In this article, I have considered the problem of separating causal from spurious components of covariances. More generally, one might be interested in estimating how each of multiple alternative causal pathways contributes to a covariance. By leveraging the chain rule of calculus, the gradient definition extends naturally to provide additive partitions of such path-specific effects (Henshaw et al. 2020; Frank 2026). This allows one to partition the causal covariance into components arising via all contributing causal pathways, analogous to Wright’s rules in path analysis (Wright 1934; Loehlin 2004). In contrast, extending the two-step definition to estimate path-specific effects is more difficult, because the analysis of finite differences naturally leads to non-additivity. One can avoid non-additivity by specifying an order in which to analyse causal pathways. However, there is no natural ordering for a set of causal pathways, and so this approach introduces arbitrary decisions with no causal basis. Alternatively, one can average over all possible orderings, analogous to the calculation of the Shapley values (Shapley 1953; Osborne and Rubinstein 1994). The method of Singal and Michailidis (2024), which is based on Shapley values, provides a particularly elegant solution. A catch, however, is that the effect attributed to a particular pathway by this method is not independent of how the remaining pathways are partitioned (e.g., a three-way contrast between paths *A, B* and *C* may result in a different estimate for the effect of *A* than a two-way contrast between *A* and the merged pathways {*B, C*}).

In conclusion, I have provided two candidate solutions to the neglected problem of determining what part of a covariance results from the causal effect of one variable on the other. Both approaches fulfil minimal requirements for a definition of causal covariance. They differ mainly in the way causal effects are decomposed and averaged – based on local vs large changes in the variables – and in their distributional assumptions. These definitions are related to similar concepts in a great variety of disciplines. In particular, they can be applied to partition the association between a trait and fitness into a causal component – which derives from the trait’s direct or indirect effects on fitness – and a spurious component arising via other pathways. In this context, the definitions operationalise Sober’s (1984) distinction between ‘selection for’ a trait, which corresponds to the causal component, and ‘selection of’ a trait, which corresponds to the total association.

## AI use

I used ChatGPT (OpenAI 2026) to refine language and find relevant literature. All ideas are my own.

## Supporting information

Data for worked example

R code for worked example

## Acknowledgments

I am grateful to Michael Morrissey, Junzhe Zhang, Larry Goldstein and Gesine Reinert for helpful discussions and assistance in navigating the literature. Funding was provided by the German Federal Ministry of Research, Technology and Space (BMFTR) and the German Research Foundation (DFG, project number 456626331).

## Appendix

### Extension of the gradient definition to non-normal variables

Here I discuss a possible extension of the gradient definition when the focal variable *X* is not normally distributed. For normal random variables *X*, Stein’s lemma links the covariance cov(*X, Z*) to the mean derivative of the conditional expectation E (*Z*|*X*) with respect to *X* (Stein 1981; Liu 1994):

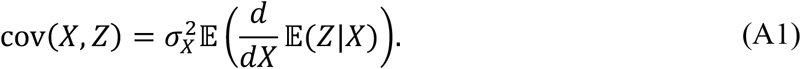

When *X* is not normal, Goldstein and Reinert (1997, 2005) showed that there exists a related random variable *X*′ such that^3^:

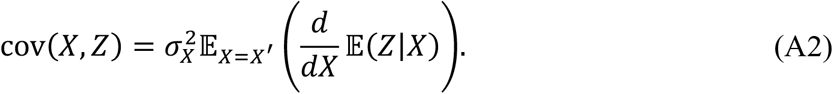

The random variable *X*′ is called the *zero-bias transform* of *X* (Figure A1). It is absolutely continuous with probability density function given by (Goldstein and Reinert 1997, 2005):

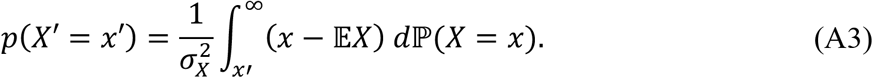

The distributions of *X* and *X*′ coincide if and only if *X* is normal. When the distribution of *X* is nearly normal, the original and zero-bias distributions are very similar, whereas large deviations from normality can result in substantial divergence between these two distributions (Figure A1). The mode of *X*′ always coincides with the mean of the original distribution. The support of *X*′ is equal to the closed convex hull of the support of *X*. Note that *X*′ depends only on the distribution of *X* and not on its relationship to any other variables.

Analogously to the gradient definition in the main text, one could conceivably define:

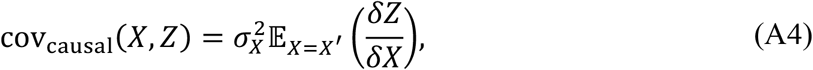

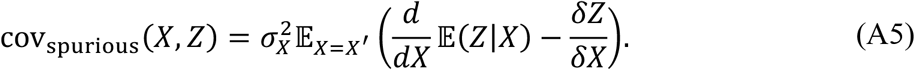

This is the *zero-bias definition* of causal and spurious covariances. It fulfils the minimal requirements (R1) through (R3) and reduces to the gradient definition when *X* is normal. However, the causal interpretation of the zero-bias definition is not straightforward, because effects are averaged over the zero-bias distribution instead of the empirical distribution. I am not aware of a causal interpretation of the zero-bias transform itself.

**Figure A1:**
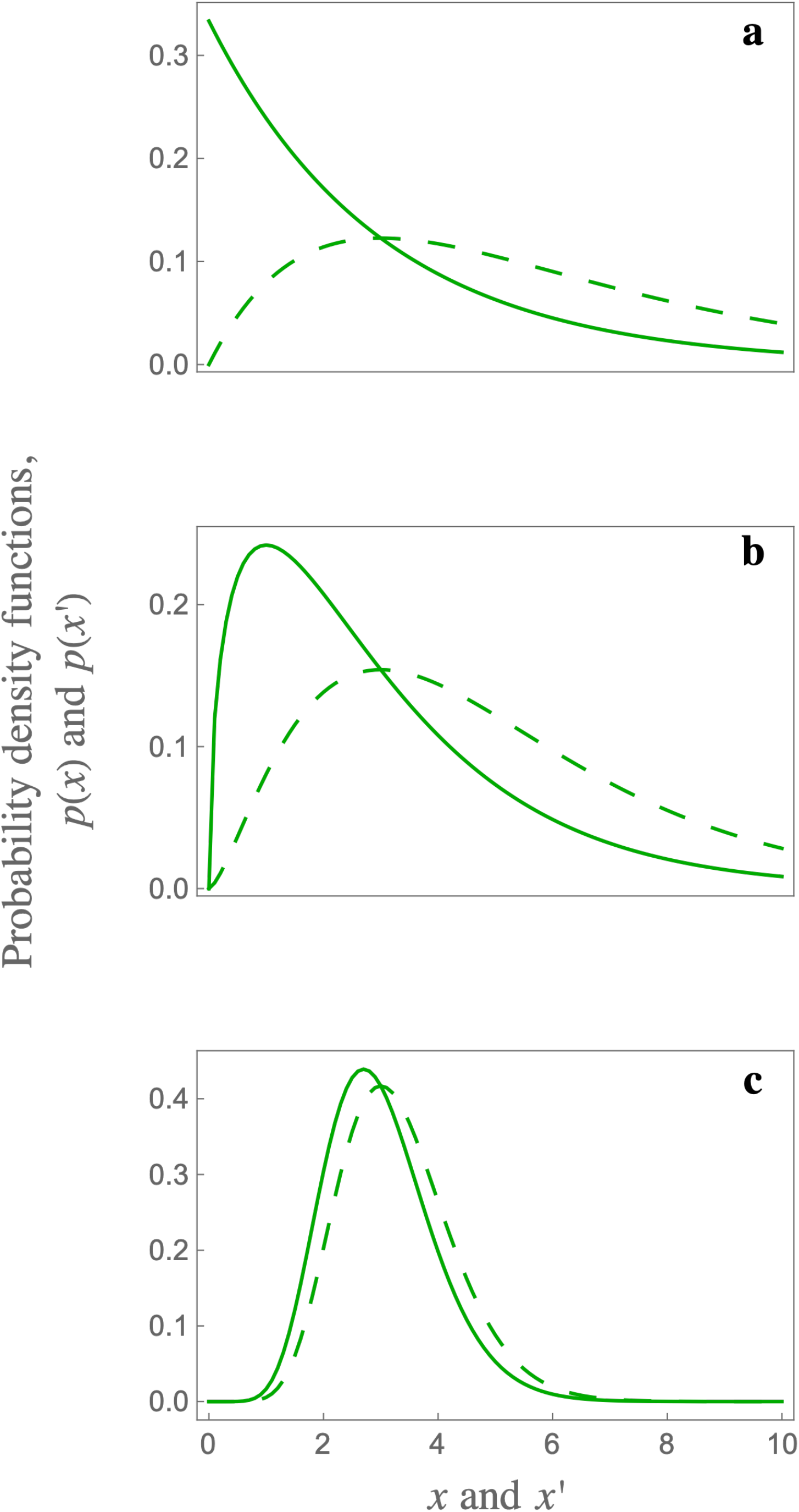
Zero-bias transforms of some common distributions. Shown are the probability density functions of the original random variable *X* (solid lines) and its zero-bias transform *X*′ (dashed lines) when *X* follows: (**a**) an exponential distribution with rate parameter 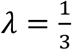 (**b**) a chi-squared distribution with 3 degrees of freedom; and (**c**) a gamma distribution with shape parameter *k* = 10 and scale parameter 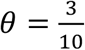. The mode of the zero-bias transform always coincides with the mean of the original distribution (Goldstein and Reinert 1997), which for the illustrated distributions is E*X* = 3. If *X* is nearly normal, then the distributions of *X* and *X*′ are similar (e.g., panel **c**). Under strong deviations from normality, the zero-bias transform differs substantially from the original distribution (e.g., panel **a**).

## Footnotes

1 A fourth possibility occurs when observations of *X* and *Z* are selected conditional on some third variable(s) and this conditioning ‘unblocks’ causal pathways between *X* and *Z*. The simplest case is *collider bias*, where *X* and *Z* both affect a third variable *C* (*X* → *C* ← *Z*). Even if *X* and *Z* are (unconditionally) independent, conditioning on *C* can induce covariance between these variables. For example, an animal might produce an alarm call in response to two different types of predators, snakes and hawks. Suppose that the presence of hawks is independent of the presence of snakes. After hearing an alarm call, I expect that either a hawk or a snake is likely present. If I then see a snake, it reduces my estimated probability that a hawk is present, and vice versa. Thus, two previously independent events (snake presence and hawk presence) become dependent conditional on a third variable (alarm call).

2 Note that (D2) corresponds to the definition of Zhang and Bareinboim (2018) if one follows the suggestion in their footnote 8.

3 Here we use the notational shorthand E_*x=x′*_*g*(*X, Z*) *≔* E_*x′*_ (*g*(*X, Z*)|_*x=x′*_) for any function *g*(*X, Z*). In other words, we take the average of *g*(*X, Z*) over the probability distribution of *X*′ rather than *X*.

