## Supplementary material for "Partitioning covariance into causal and spurious components": R code for worked example

# This document provides R code
# for the worked example in:

# Partitioning covariance into
# causal and spurious components

# This code was written in R v4.4.3.

# Set-up ------------------------------------------------------------------

# load packages
library(ggplot2)
library(emmeans)

# import the data set
Pipefish <- read.csv("PipefishData.csv")

# regression model to predict egg number
# in terms of body size, collection event (= site)
# and their interaction
model <- lm(eggs ~ size * site, data = Pipefish)
summary(model)

# plot the regression model against the data

okabe_ito <- c("#E69F00", "#56B4E9", "#009E73",
               "#F0E442", "#0072B2", "#D55E00",
               "#CC79A7", "#000000")

ggplot(Pipefish, aes(x=size, y=eggs, color=site)) +
  geom_point() + 
  geom_smooth(method="lm", se=F) +
  theme_minimal() +
  labs(x = expression("body size, " * italic("S") * " (mm)"),
       y = expression("number of eggs, "*italic("E")),
       color = expression("collection event, "*italic("C"))) +
  scale_colour_manual(values = okabe_ito)

# calculate total covariance cov(E,S)
cov_true <- cov(Pipefish$size, Pipefish$eggs)

# Gradient definition -----------------------------------------------------

# I provide two approaches for calculating
# the causal and spurious covariance
# of body size with egg number.

# These two approaches are essentially
# identical in output.

## emtrends() approach -----------------------------------------------------

emtrends_model <- emtrends(
  model,
  specs = ~ 1,
  var = "size",
  data = Pipefish,
  cov.reduce = FALSE,
  weights = "proportional"
  )

summary(emtrends_model)

# extract the mean δE/δS from the emtrends model
mean_gradient_emtrends <- summary(emtrends_model)$size.trend

# calculate cov_causal(E,S)
# using the gradient definition
cov_causal_gradient_emtrends <- mean_gradient_emtrends*var(Pipefish$size)

# cov_spurious(E,S) is the remainder
cov_spurious_gradient_emtrends <- cov_true - cov_causal_gradient_emtrends


## manual numeric approach -------------------------------------------------

# magnitude of small change for
# calculating gradients numerically
epsilon <- 0.1

# data set with all size values increased by ε
Pipefish_epsilon <- Pipefish %>%
  mutate(size = size + epsilon)

# mean δE/δS
mean_gradient_gradient_manual <- 1/epsilon * mean(predict(model, Pipefish_epsilon, type = "response") - predict(model, Pipefish, type = "response"))

# calculate cov_causal(E,S)
# using the gradient definition
cov_causal_gradient_manual <- mean_gradient_manual*var(Pipefish$size)

# cov_spurious(E,S) is the remainder
cov_spurious_gradient_manual <- cov_true - cov_causal_gradient_manual

# Two-step definition -----------------------------------------------------

# set seed for reproducibility
set.seed(2025)

# number of times to simulate paired draws
# from the joint distribution of variables
num_samples <- 10^6

# random indices for resampling
indices <- sample(nrow(Pipefish), num_samples, replace = T)
indices_star <- sample(nrow(Pipefish), num_samples, replace = T)

# resamples (S,C,E)
samples <- Pipefish[indices,]

# resamples (S*,C,E)
samples_star <- samples %>%
  mutate(size = Pipefish$size[indices_star])

# S
S <- samples$size

# S*
S_star <- samples_star$size

# extract residual variance from regression model
sigma <- summary(model)$sigma

# simulate E|S,C
E <- predict(model, samples, type = "response") + rnorm(num_samples, mean = 0, sd = sigma)

# simulate E|S*,C
E_do_S_star <- predict(model, samples_star, type = "response") + rnorm(num_samples, mean = 0, sd = sigma)

# calculate cov(E,S)
# using re-sampled S and model-simulated E|S,C
cov_true_twostep <- cov(S, E)

# calculate cov_causal(E,S)
# using the two-step definition
cov_causal_twostep <- 0.5*(cov(S_star, E_do_S_star) +
                             cov(S, E) -
                             cov(S, E_do_S_star))

# calculate cov_spurious(E,S)
# using the two-step definition
cov_spurious_twostep <- 0.5*(cov(S, E) -
                               cov(S_star, E_do_S_star) +
                               cov(S, E_do_S_star))
